# Seed Amplification Assay–Based Detection of Alpha-Synuclein Aggregates Outperforms Immunohistochemical Analysis in the Intestinal Mucosal Biopsies of Parkinson’s Disease and Multiple System Atrophy Participants

**DOI:** 10.64898/2026.11.05.764662

**Authors:** Ali Keshavarzian, Phillip A. Engen, Claudio Soto, Robin M. Voigt, Bryan Killinger, Maliha Shaikh, Michelle Pinho, Sandra Pritzkow, Jayda Duvernay, Mariah Barnett, Stefan J. Green, Mohamed Abdel-Mohsen, Amol Sharma, Christopher G. Goetz

## Abstract

**Objective:** To determine whether an α-synuclein (α-Syn) seed-amplification-assay (SAA) detects seeding-competent aggregates in endoscopic mucosal biopsies, differs by gastrointestinal region, registers pathology more frequently than immunohistochemistry (IHC), and associates with Parkinson’s disease (PD) diagnosis, disease duration or severity.

**Design:** This cross-sectional, proof-of-concept diagnostic accuracy study analyzed duodenum, ileum, and sigmoid biopsies from PD participants, age- and sex-matched healthy controls (HC), plus exploratory multiple system atrophy (MSA) subtypes [parkinsonian (MSA-P) and cerebellar (MSA-C)]. Ileal and sigmoid PD and HC biopsies were assessed using proteinase K-resistant phospho-Ser129 IHC. SAA-positivity was evaluated against clinical and biological measures.

**Results:** In PD, SAA detected seeding-competent α-Syn aggregates with high observed specificity (97.5–100%) and region-dependent sensitivity (18/37, 48.65% in ileum and 36/56, 64.29% in sigmoid). Sigmoid SAA positivity increased significantly with PD disease duration (40.00% at <5 years, 73.33% at 5–10 years, and 93.75% at >10 years), independent of motor severity, whereas ileal positivity remained stable across disease stages. In 37 paired sigmoid biopsies, SAA detected α-Syn aggregates 2.8-fold more frequently than IHC. Sigmoid seeding activity was detected in all five MSA-P, but not in two MSA-C. IHC did not detect aggregates in any MSA.

**Conclusions:** In PD, α-Syn SAA detected seeding-competent α-Syn-aggregates in endoscopic mucosal biopsies with high observed specificity, moderate sensitivity, and approximately 2.8-fold higher detection rate than IHC in sigmoid. Positive SAA in MSA patients supports GI involvement across synucleinopathies. SAA detection in PD differed by intestinal region and disease duration, supporting further validation of intestinal SAA as a PD biomarker.

**Significance of this study:** *What is already known about this subject?:*

- Alpha-synuclein (α-syn) seed-amplification-assay (SAA) is highly accurate in cerebrospinal fluid and skin, but collection requires a dedicated invasive or minor surgical procedure and is not collected at population scale.
- Four previous studies have reported α-syn seeding activity in gastrointestinal biopsies, with sample sizes of 7–23 patients, single anatomical regions, and widely varying positivity rates.
- Immunohistochemistry for α-syn in mucosal biopsies is limited by sparse, unevenly distributed pathology, the absence of a standardized scoring system, and its inability to establish whether detected aggregates are seeding-competent.
- Whether gastrointestinal seeding activity varies by anatomical region, disease duration, disease severity or severity of motor and GI symptoms is unknown.

*What are the new findings?:*

- In the largest gastrointestinal α-Syn SAA series to date (96 participants), biopsies obtained during routine endoscopy identified PD with 97.50-100% specificity and moderate sensitivity (48.65% ileum, 64.29% sigmoid).
- Sigmoid SAA-positivity was duration dependent, rising from 40% at <5 years to 94% beyond 10 years (adjusted OR 2.82 per 5 years), and disease duration was the only independent correlate once H&Y stage was included in the model.
- Constipation, intestinal inflammation, and sigmoid mucosal dysbiosis tracked PD severity but did not distinguish sigmoid SAA-positive from SAA-negative participants.
- Ileal SAA-positivity was constant across disease duration and severity, indicating that positivity was not uniform along the gastrointestinal tract.
- In the identical PD participant’s sigmoid tissue, SAA identified α-Syn pathology in 28 of 37 patients compared with 10 of 37 by proteinase K-resistant phospho-Ser129 immunohistochemistry.
- Sigmoid SAA-positivity was detected in 5 of 7 participants with MSA, with positive cases confined to the parkinsonian subtype (MSA-P) and negative cases to the cerebellar subtype (MSA-C).
- Strain typing by α-Syn SAA enables differentiation of PD from MSA cases using sigmoid biopsies.

*How might it impact on clinical practice in the foreseeable future?:*

- Colonoscopy is already performed at scale in the age group at-risk of PD, so mucosal biopsies are an accessible matrix for α-Syn seeding assays and for biomarker-based enrichment of clinical trials.
- Because yield depends on both sampling site and disease duration, prospective validation against cerebrospinal fluid must specify region and stage; a negative result from a single site cannot currently exclude PD.

## Introduction

Parkinson’s disease (PD) is among the most common and fastest-growing neurodegenerative disorders(1), yet the diagnosis of PD remains clinical, and disease-modifying trials still lack objective, accessible markers of alpha-synuclein (α-syn) pathology(2). Seed-amplification-assays (SAA) exploit the capacity of pathological α-syn to template the misfolding and aggregation of recombinant α-syn, achieving high diagnostic performance in cerebrospinal fluid (CSF), with reported SAA sensitivity of 88-92% and specificity of 95-97% for Lewy body synucleinopathies(3). CSF sampling is invasive and not readily repeatable at scale, which has driven interest in peripheral matrices including skin and the gastrointestinal (GI) tract(4, 5).

The GI tract is a particularly appealing matrix because mucosal biopsies are already obtained during routine colonoscopy in the age group at risk of PD, and because α-syn aggregates are present in colonic tissue years before motor symptom onset(5–7). Immunohistochemistry (IHC), however, has repeatedly failed as a diagnostic readout in mucosal biopsies primarily because pathology is sparse, unevenly distributed, and aggregate specific antibodies are often immunoreactive in healthy controls (HC), and staining cannot establish if what is detected is seeding-competent(8–13).

Four prior studies have used seeding assays in GI biopsies. Fenyi et al. detected α-syn aggregates by protein misfolding cyclic amplification in sigmoid biopsies from 58.33% of (7/12) PD patients(14), Shin et al. reported SAA-positivity in gastric tissue from 10.00% of (2/20) PD patients(15), Vascellari et al. found seeding activity in duodenal biopsies from 95.65% of (22/23) PD patients(16), and Emmi et al. report SAA-positivity in 40.91% of duodenum of (9/22) PD patients(17). These reports establish feasibility but were small in sample size, limited in comparison between anatomical regions of the GI tract, did not compare SAA with IHC in the same tissue, did not assess if α-syn-positivity varied with disease duration, disease severity, a biomarker of neurodegeneration, symptoms severity, and mucosal-associated microbiota community.

This study used α-Syn SAA and proteinase K-resistant, phospho-Ser129 (pS129) IHC to assess intestinal mucosal biopsies (duodenum, ileum, sigmoid) from participants with PD and HC to determine if intestinal seeding-competent α-Syn aggregates can (1) distinguish PD from HC, (2) vary by anatomical GI region, and (3) track disease duration, clinical severity, and correlate with biomarkers of neurodegeneration, symptom burden, mucosal dysbiosis, or intestinal inflammation. Exploratory analysis also evaluated SAA-positivity in sigmoid biopsies from seven participants with multiple system atrophy (MSA), another central nervous system synucleinopathy, with two classic phenotypes, parkinsonism (MSA-P) and cerebellar ataxia (MSA-C).

## Methods

### Participants

The Rush University Medical Center Institutional Review Board approved the study (ORA: 23102401), and participants provided written informed consent. A movement disorders specialist (CGG) confirmed PD and MSA diagnoses according to diagnostic criteria(18, 19). Motor severity was assessed using the MDS-Unified Parkinson’s Disease Rating Scale (MDS-UPDRS) Part III(20), and Hoehn and Yahr (H&Y) staging (21). PD patients were divided into three severity groups, defined operationally by the absence (H&Y: 1-2) or presence (H&Y: 3,4,5) of balance problems and dopaminergic medication use: mild: H&Y 1-2 and no medication; moderate H&Y 1-2 but on medication; severe H&Y ≥ 3 and on medication use. Healthy controls (HC) were age-, sex-, and race-matched to PD participants. Participant demographics and clinical characteristics are depicted in **Table 1** and **Source Data**. The seven MSA participants (n=2 cerebellar subtype (MSA-C), n=5 parkinsonian subtype (MSA-P)) demographics are shown in **Table 2** and **Source Data**.

**Table 1.**
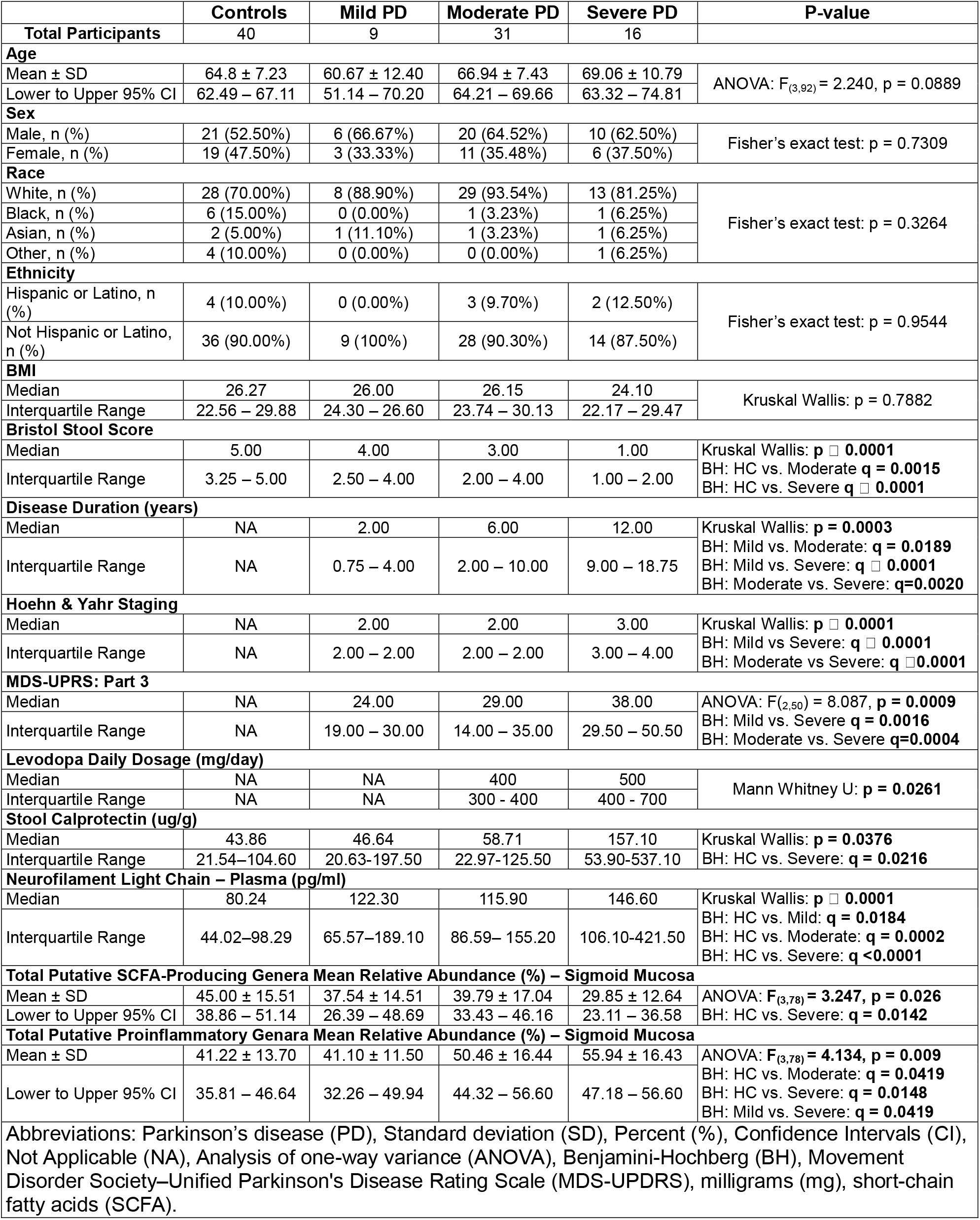
Demographics, Clinical, and Experimental Assessments.

|  | **Controls** | **Mild PD** | **Moderate PD** | **Severe PD** | **P-value** |
|---|---|---|---|---|---|
| **Total Participants** | 40 | 9 | 31 | 16 |  |
| **Age** |  |  |  |  |  |
| Mean ± SD | 64.8 ± 7.23 | 60.67 ± 12.40 | 66.94 ± 7.43 | 69.06 ± 10.79 | ANOVA: F<sub>(3,92)</sub> = 2.240, p = 0.0889 |
| Lower to Upper 95% CI | 62.49 – 67.11 | 51.14 – 70.20 | 64.21 – 69.66 | 63.32 – 74.81 |  |
| **Sex** |  |  |  |  |  |
| Male, n (%) | 21 (52.50%) | 6 (66.67%) | 20 (64.52%) | 10 (62.50%) | Fisher’s exact test: p = 0.7309 |
| Female, n (%) | 19 (47.50%) | 3 (33.33%) | 11 (35.48%) | 6 (37.50%) |  |
| **Race** |  |  |  |  |  |
| White, n (%) | 28 (70.00%) | 8 (88.90%) | 29 (93.54%) | 13 (81.25%) | Fisher’s exact test: p = 0.3264 |
| Black, n (%) | 6 (15.00%) | 0 (0.00%) | 1 (3.23%) | 1 (6.25%) |  |
| Asian, n (%) | 2 (5.00%) | 1 (11.10%) | 1 (3.23%) | 1 (6.25%) |  |
| Other, n (%) | 4 (10.00%) | 0 (0.00%) | 0 (0.00%) | 1 (6.25%) |  |
| **Ethnicity** |  |  |  |  |  |
| Hispanic or Latino, n (%) | 4 (10.00%) | 0 (0.00%) | 3 (9.70%) | 2 (12.50%) | Fisher’s exact test: p = 0.9544 |
| Not Hispanic or Latino, n (%) | 36 (90.00%) | 9 (100%) | 28 (90.30%) | 14 (87.50%) |  |
| **BMI** |  |  |  |  |  |
| Median | 26.27 | 26.00 | 26.15 | 24.10 | Kruskal Wallis: p = 0.7882 |
| Interquartile Range | 22.56 – 29.88 | 24.30 – 26.60 | 23.74 – 30.13 | 22.17 – 29.47 |  |
| **Bristol Stool Score** |  |  |  |  |  |
| Median | 5.00 | 4.00 | 3.00 | 1.00 | Kruskal Wallis: **p □ 0.0001**<br>BH: HC vs. Moderate **q = 0.0015**<br>BH: HC vs. Severe **q □ 0.0001** |
| Interquartile Range | 3.25 – 5.00 | 2.50 – 4.00 | 2.00 – 4.00 | 1.00 – 2.00 |  |
| **Disease Duration (years)** |  |  |  |  |  |
| Median | NA | 2.00 | 6.00 | 12.00 | Kruskal Wallis: **p = 0.0003**<br>BH: Mild vs. Moderate: **q = 0.0189**<br>BH: Mild vs. Severe: **q □ 0.0001**<br>BH: Moderate vs. Severe: **q=0.0020** |
| Interquartile Range | NA | 0.75 – 4.00 | 2.00 – 10.00 | 9.00 – 18.75 |  |
| **Hoehn & Yahr Staging** |  |  |  |  |  |
| Median | NA | 2.00 | 2.00 | 3.00 | Kruskal Wallis: **p □ 0.0001**<br>BH: Mild vs Severe: **q □ 0.0001**<br>BH: Moderate vs Severe: **q □0.0001** |
| Interquartile Range | NA | 2.00 – 2.00 | 2.00 – 2.00 | 3.00 – 4.00 |  |
| **MDS-UPRS: Part 3** |  |  |  |  |  |
| Median | NA | 24.00 | 29.00 | 38.00 | ANOVA: F(<sub>2,50</sub>) = 8.087, **p = 0.0009**<br>BH: Mild vs. Severe **q = 0.0016**<br>BH: Moderate vs. Severe **q=0.0004** |
| Interquartile Range | NA | 19.00 – 30.00 | 14.00 – 35.00 | 29.50 – 50.50 |  |
| **Levodopa Daily Dosage (mg/day)** |  |  |  |  |  |
| Median | NA | NA | 400 | 500 | Mann Whitney U: **p = 0.0261** |
| Interquartile Range | NA | NA | 300 - 400 | 400 - 700 |  |
| **Stool Calprotectin (ug/g)** |  |  |  |  |  |
| Median | 43.86 | 46.64 | 58.71 | 157.10 | Kruskal Wallis: **p = 0.0376** |
| Interquartile Range | 21.54–104.60 | 20.63-197.50 | 22.97-125.50 | 53.90-537.10 | BH: HC vs. Severe: **q = 0.0216** |
| **Neurofilament Light Chain – Plasma (pg/ml)** |  |  |  |  |  |
| Median | 80.24 | 122.30 | 115.90 | 146.60 | Kruskal Wallis: **p □ 0.0001**<br>BH: HC vs. Mild: **q = 0.0184**<br>BH: HC vs. Moderate: **q = 0.0002**<br>BH: HC vs. Severe: **q <0.0001** |
| Interquartile Range | 44.02–98.29 | 65.57–189.10 | 86.59– 155.20 | 106.10-421.50 |  |
| **Total Putative SCFA-Producing Genera Mean Relative Abundance (%) – Sigmoid Mucosa** |  |  |  |  |  |
| Mean ± SD | 45.00 ± 15.51 | 37.54 ± 14.51 | 39.79 ± 17.04 | 29.85 ± 12.64 | ANOVA: **F<sub>(3,78)</sub> = 3.247, p = 0.026** |
| Lower to Upper 95% CI | 38.86 – 51.14 | 26.39 – 48.69 | 33.43 – 46.16 | 23.11 – 36.58 | BH: HC vs. Severe: **q = 0.0142** |
| **Total Putative Proinflammatory Genara Mean Relative Abundance (%) – Sigmoid Mucosa** |  |  |  |  |  |
| Mean ± SD | 41.22 ± 13.70 | 41.10 ± 11.50 | 50.46 ± 16.44 | 55.94 ± 16.43 | ANOVA: **F<sub>(3,78)</sub> = 4.134, p = 0.009** |
| Lower to Upper 95% CI | 35.81 – 46.64 | 32.26 – 49.94 | 44.32 – 56.60 | 47.18 – 56.60 | BH: HC vs. Moderate: **q = 0.0419**<br>BH: HC vs. Severe: **q = 0.0148**<br>BH: Mild vs. Severe: **q = 0.0419** |

Abbreviations: Parkinson’s disease (PD), Standard deviation (SD), Percent (%), Confidence Intervals (CI), Not Applicable (NA), Analysis of one-way variance (ANOVA), Benjamini-Hochberg (BH), Movement Disorder Society–Unified Parkinson's Disease Rating Scale (MDS-UPDRS), milligrams (mg), short-chain fatty acids (SCFA).

**Table 2.** Demographics and Clinical Assessment of the Multiple System Atrophy Participants.

| **Total Participants**, n (%) | **n=7**<br>n=2 (28.57%), MSA-C<br>n=5 (71.43%), MSA-P |
|---|---|
| **Age** |  |
| Mean ± SD | 69.29 ± 6.921 |
| Lower to Upper 95% CI | 62.88 – 75.69 |
| **Sex** |  |
| Male, n (%) | 6 (85.71%) |
| Female, n (%) | 1 (14.29%) |
| **Race** |  |
| White, n (%) | 7 (100%) |
| **Ethnicity** |  |
| Not Hispanic or Latino, n (%) | 7 (100%) |
| **BMI** |  |
| Mean ± SD | 30.06 ± 4.256 |
| Lower to Upper 95% CI | 26.12 – 33.99 |
| **Disease Duration (years)** |  |
| Median | 6.00 |
| Interquartile Range | 3.00 – 9.00 |
| **Hoehn and Yahr Staging** |  |
| Median | 3.00 |
| Interquartile Range | 3.00 - 4.00 |
| **MDS-UPDRS: Part 3** |  |
| Median | 38.20 |
| Interquartile Range | 31.10 – 49.00 |
| **Constipation Duration (years)** |  |
| Mean ± SD | 3.38 ± 1.03 |
| **Constipation Severity Score** |  |
| Mean ± SD | 3.83 ± 1.33 |
| **Stool Calprotectin (ug/g)** |  |
| Median | 168.8 |
| Interquartile Range | 49.54 – 424.00 |

**MSA-P**: Multiple system atrophy, parkinsonian type, characterized predominantly by parkinsonism, including slowness of movement, rigidity, and postural instability.
**MSA-C:** Multiple system atrophy, cerebellar type, characterized predominantly by cerebellar dysfunction, including impaired balance, coordination, and speech.
Medication usage.
Antiparkinsonian medication data were unavailable for the MSA participants and therefore could not be reported.
Abbreviations: Standard deviation (SD), Percent (%), Confidence Intervals (CI), Movement Disorder Society–Unified Parkinson's Disease Rating Scale (MDS-UPDRS), MSA (Multiple System Atrophy).

### Blood and Stool Collection

Blood was collected in K2-EDTA tubes (BD #366643) by a trained phlebotomist, processed (800 x g; 15 min; 9 acceleration; 9 brake) within one-hour of collection, and plasma was aliquoted and stored at −80°C.

Participants collected stool at home within 24 h of the study visit, using an anaerobic home collection kit (BD GasPak:260683, Becton Dickinson and Company, Sparks, MD). Participants recorded the Bristol Stool Score (types 1-2 = constipation, types 3-4 = normal stool, types 5-7 loose stool/diarrhea) which was verified by a clinical coordinator. Samples were stored at −80°C until use.

### Biopsy Collection

Four-millimeter mucosa biopsies were obtained from the ileum and sigmoid colon during clinically indicated colonoscopy, and from the duodenum during clinically indicated upper gastrointestinal endoscopy (EGD). Individual biopsy wet weights were approximately 5–20 mg. Only participants with normal-appearing mucosa (defined by an experienced endoscopist-AK) were included in the study. Biopsies were placed into sterile cryovials, flash-frozen in liquid nitrogen, and stored at −80°C.

### Sigmoid Mucosa Microbiota

DNA was extracted from sigmoid mucosa biopsies using the Qiagen DNeasy PowerSoil kit. Near full-length 16S rRNA genes were amplified and sequenced on a PacBio Revio instrument(22). Reads were processed in QIIME2 for quality control, DADA2 ASV generation, and taxonomic assignment using GTDB r226(23, 24). Genera were categorized as putative short-chain fatty acid-producing taxa or pro-inflammatory taxa(25), and their relative abundances were summed to calculate total abundance values and compared across groups(26–29).

### Blood and Stool Assay

Plasma neurofilament light chain (NfL), a systemic biomarker of neuroaxonal injury and neurodegeneration(30), was measured using the R-PLEX Human Neurofilament L Assay (Meso Scale Discovery; K1517XR-2) according to the manufacturer’s instructions. Stool calprotectin, a biomarker of intestinal inflammation(31), was measured using the BÜHLMANN Calprotectin ELISA kit (EK-CAL2) according to the manufacturer’s instructions.

### Recombinant α-Synuclein Preparation

Recombinant human α-Syn was produced and purified using an established protocol with modifications(32, 33), as detailed in **Supplementary Methods**.

### Alpha-Synuclein Seed Amplification Assay

The α-Syn SAA was used to detect seeding-competent α-Syn aggregates in intestinal mucosal biopsies using a previously described protocol(34, 35). Biopsies collected from the duodenum, ileum, and sigmoid colon were processed to generate tissue homogenates for SAA analysis. Tissue homogenate was diluted 10000-fold to remove interferences with SAA. For each reaction, 40 µL of diluted tissue homogenate was combined with 60 µL of reaction solution in a clear-bottom 96-well plate (Greiner). The reaction solution contained 0.2 mg/ml recombinant 6×His-tagged αSyn, 100 mM PIPES (pH 6.5; Sigma-Aldrich), 500 mM NaCl, and 10 µM thioflavin T (ThT). Each reaction well also contained two 1/8-inch silicon nitride beads (Xiamen MasCera Technology) to facilitate aggregate amplification.

Plates were incubated at 42 °C in an Omega microplate reader. Samples underwent orbital agitation at 800 rpm for 1 min at 14-min intervals, with ThT fluorescence recorded following each shaking cycle. All biopsy specimens were evaluated in triplicate, with amplification monitored for 24 h.

A fluorescence value greater than 1,000 arbitrary units (AU) was used as the predetermined positivity threshold. Samples were classified as SAA-positive when at least two of the three technical replicates exceeded this threshold. Samples with zero or one positive replicate were classified as negative. As previously reported, the maximum fluorescence in the SAA reaction enables the test to distinguish patients affected by PD from MSA(32, 37). When the fluorescence signal of at least two of the three replicates is between 2000 and 25000, the samples are classified as having the MSA strain pattern; whereas when the maximum fluorescence of at least two of the three replicates is >25000, the sample contains the Lewy body disease (LBD) strain pattern.

### Immunohistochemistry

Formalin-fixed paraffin-embedded biopsies were treated with proteinase K and immunostained for phosphorylated α-synuclein (pS129; EP1536Y, Abcam)(36–38). Two to four slides per biopsy were independently scored by three blinded reviewers using a 0-3 scale(39). Detailed staining procedures are provided in **Supplementary Methods**.

### Statistics

Analyses were performed in GraphPad Prism 10 (San Diego, CA). Distributional assumptions were evaluated with the Shapiro–Wilk test. Two-group comparisons used the Mann–Whitney U test, and multiple-group comparisons used one-way ANOVA or the Kruskal–Wallis test, as appropriate; specified post-hoc comparisons were adjusted by the Benjamini–Hochberg (BH) procedure. Two-way ANOVA assessed PD group, sigmoid SAA detection status, and their interaction for continuous outcomes. Categorical outcomes were compared with Fisher’s exact test. Paired SAA–IHC detection was compared using McNemar’s exact test. Logistic regression was used to examine associations between disease duration, clinical measures, and sigmoid SAA-positivity. Nominal p<0.05 was considered statistically significant for unadjusted comparisons; where BH correction was applied, q<0.05 was used. Counts, detection proportions, sensitivity, specificity, and case-control sample-based predictive values were calculated.

## Results

Participant demographics and clinical characteristics across the three operational PD subgroups are presented in **Table 1**. The three PD severity categories behaved as expected: disease duration increased mild, moderate, and severe (median 2, 6, and 12 years, respectively; KW: p=0.0003) and MDS-UPDRS: Part III scores were higher with more advanced disease (ANOVA: F_(2,50)_ = 8.087, p=0.0009). Age, sex, race, ethnicity, and body mass index did not differ significantly across groups (**Table 1**). Subsequent analyses addressed associations of intestinal seeding-competent α-Syn aggregates with clinical disease features and with the measured gastrointestinal phenotype.

### Intestinal SAA Detects Seeding Competent α-Synuclein Aggregates in PD and an Exploratory MSA Cohort

SAA was performed on ileal biopsies from 36 HC and 37 PD participants (2 mild, 25 moderate, 10 severe); sigmoid biopsies from 40 HC and 56 PD participants (9 mild, 31 moderate, 16 severe); and duodenal biopsies from 1 HC and 7 PD participants (0 mild, 4 moderate, 3 severe). Seeding competent α-syn aggregates were positively detected in 48.65% of ileum (18/37), 64.29% of sigmoid (36/56), and 71.43% of duodenum (5/7) PD biopsies (**Table 3**, **Figure 1A-C**). Among HC GI biopsies, 0/36 ileal and 1/40 sigmoid biopsies were positive, corresponding to observed specificities of 100% and 97.50%, respectively. The duodenum comparison included a single negative HC and is considered an exploratory analysis.

**Figure 1.**
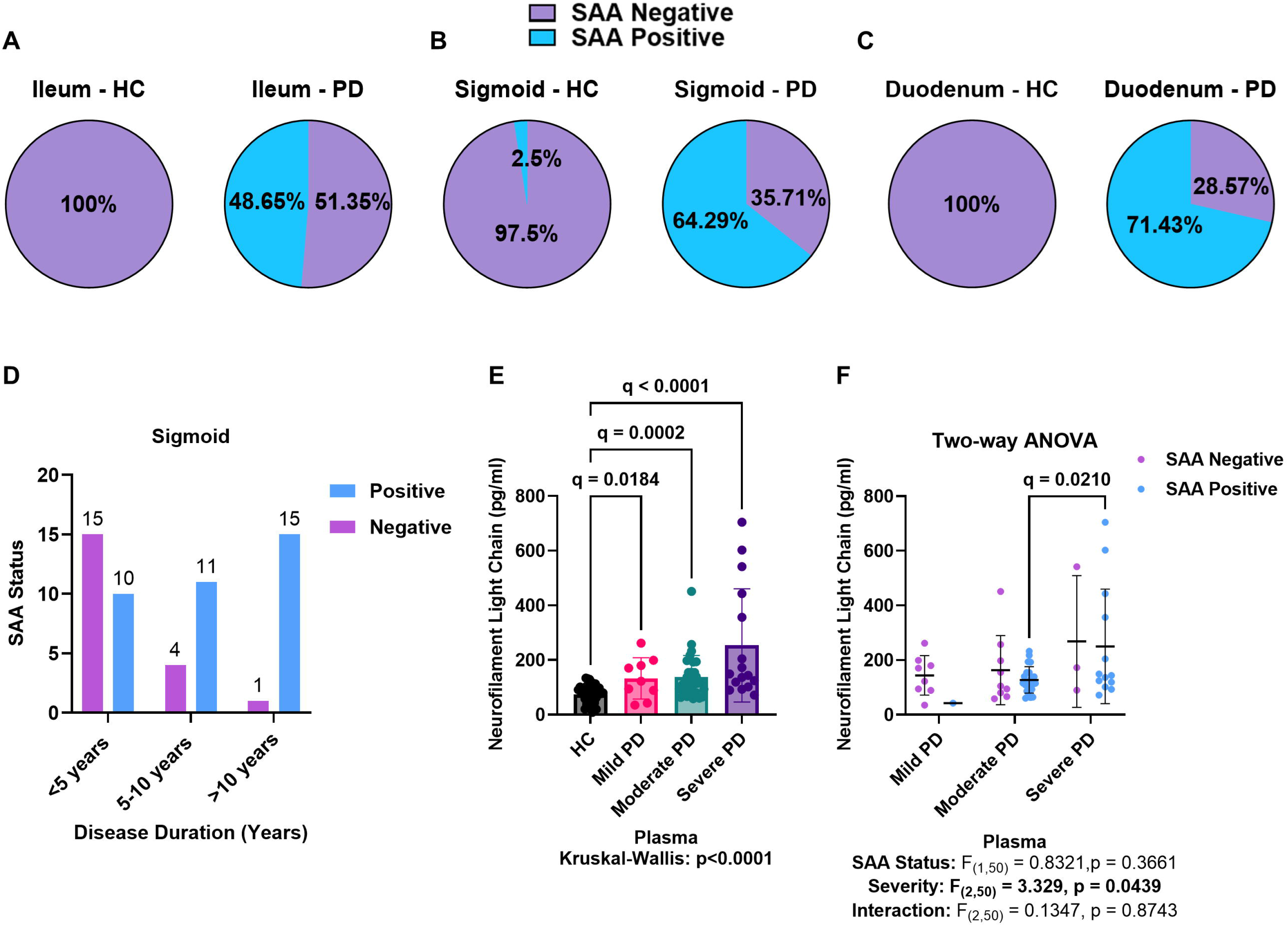
Intestinal α-synuclein SAA detection and clinical associations in Parkinson’s disease (PD). SAA-positivity in ileal (**A**), sigmoid (**B**), and duodenal (**C**) biopsies from healthy controls (HC) and PD participants. SAA-positivity were 18/37 (48.65%), 36/56 (64.29%), and 5/7 (71.43%) in PD and 0/36, 1/40, and 0/1 in HC, respectively. **(D)** Counts of sigmoid SAA-positive and SAA-negative PD participants within disease-duration categories: 10/25 (40.0%) at <5 years, 11/15 (73.3%) at 5–10 years, and 15/16 (93.8%) at >10 years. **(E)** Plasma neurofilament light chain (NfL) concentrations across HC and PD severities (Kruskal–Wallis p<0.0001); specified BH-adjusted comparisons are indicated. **(F)** Plasma NfL in PD severities stratified by sigmoid SAA status. Two-way ANOVA showed a PD severity effect (p=0.0439) but no SAA status effect (p=0.3661) or interaction (p=0.8743).

**Table 3.** Intestinal seed-amplification-assay (SAA) detection rates and diagnostic performance by biopsy site, participant groups (HC, PD, MSA), and Parkinson’s disease severity.

|  | SAA Positive, n (%) | SAA Negative, n (%) | Total | Sensitivity (%) | Specificity (%) | PPV (%) | NPV (%) |
|---|---|---|---|---|---|---|---|
| **Biopsy, Group** |  |  |  |  |  |  |  |
| Ileum, HC | 0 (0%) | 36 (100%) | 36 | - | - | - | - |
| Ileum, PD | 18 (48.65%) | 19 (51.35%) | 37 | 48.65% | 100.00% | 100.00% | 65.45% |
| Sigmoid, HC | 1 (2.50%) | 39 (97.5%) | 40 | - | - | - | - |
| Sigmoid, PD | 36 (64.29%) | 20 (35.71%) | 56 | 64.29% | 97.50% | 97.30% | 66.10% |
| Duodenal, HC* | 0 (0%) | 1 (100%) | 1 | - | - | - | - |
| Duodenal, PD* | 5 (71.43%) | 2 (28.57%) | 7 | 71.43% | 100.00% | 100.00% | 33.33% |
| Sigmoid, MSA* | 5 (71.43%) | 2 (28.57%) | 7 | 71.43% | 97.50% | 83.33% | 95.12% |
| **Biopsy, Severity** |  |  |  |  |  |  |  |
| Ileum, Mild PD | 1 (50%) | 1 (50%) | 2 | 50.00% | 100.00% | 100.00% | 97.30% |
| Ileum, Moderate PD | 12 (48%) | 13 (52%) | 25 | 48.00% | 100.00% | 100.00% | 73.47% |
| Ileum, Severe PD | 5 (50%) | 5 (50%) | 10 | 50.00% | 100.00% | 100.00% | 87.80% |
| Sigmoid, Mild PD | 1 (11.11%) | 8 (88.89%) | 9 | 11.11% | 97.50% | 50.00% | 82.98% |
| Sigmoid, Moderate PD | 22 (70.97%) | 9 (29.03%) | 31 | 70.97% | 97.50% | 95.65% | 81.25% |
| Sigmoid, Severe PD | 13 (81.25%) | 3 (18.75%) | 16 | 81.25% | 97.50% | 92.86% | 92.86% |
| Duodenum, Mild PD* | - | - | 0 | - | - | - | - |
| Duodenum, Moderate PD* | 4 (100%) | 0 (0%) | 4 | 100.00% | 100.00% | 100.00% | 100.00% |
| Duodenum, Severe PD* | 1 (33.33%) | 2 (66.67%) | 3 | 33.33% | 100.00% | 100.00% | 33.33% |

Clinical diagnosis is the reference standard. Each PD, PD severity, or MSA group is compared with the HC group from the same biopsy site; the same HC counts are reused for severity-specific estimates. HC rows do not have separate diagnostic-performance estimates. \* Exploratory analyses: duodenum HC, PD (mild, moderate and severe); sigmoid MSA. The dash (-) means not applicable; HC, healthy controls; PD, Parkinson’s disease, MSA, Multiple System Atrophy, ppv, positive predictive value; npv, negative predictive value. PPV and NPV reflect the case and control proportions in each comparison and may not generalize to populations with different disease prevalence.

SAA-positivity differed by GI region in PD participants. Ileum SAA-positivity was essentially similar across PD severity groups (mild 1/2, 50.00%; moderate 12/25, 48.00%; severe 5/10, 50.00%), although the mild PD estimate relies on only two participants. By contrast, sigmoid SAA-positivity increased with severity from 11.11% (1/9) in mild to 70.97% (22/31) in moderate and 81.25% (13/16) in severe PD (**Table 3**). Notably, the same assay used in two GI regions of the same PD participants yielded different results.

MSA was included as an additional movement disorder group for exploratory analysis. Sigmoid biopsies were SAA-positive in 5 of 7 participants (71.43%) (**Table 3**). All five SAA-positive participants had MSA-P, with disease durations of 4–9 years, whereas both SAA-negative participants had MSA-C, with shorter disease durations of 1–3 years. No region-specific seeding positivity could be assessed in MSA because these patients were too frail to have colonoscopy and underwent sigmoidoscopy.

SAA-positive MSA biopsies exhibited significantly lower maximum fluorescence values than SAA-positive PD biopsies (**Figure 2A**). Four of the five positive MSA biopsies (80.00%) exhibited an MSA-associated Type 2 seeding profile, whereas one (20.00%) exhibited a Lewy body disease (LBD)-associated Type 1 profile (**Figure 2B–C**)(35, 40). Conversely, 32 of the 36 SAA-positive PD biopsies (88.89%) exhibited an LBD-associated Type 1 seeding profile, whereas four (11.11%) exhibited an MSA-associated Type 2 profile (**Figure 2B–C**). These findings suggest that most SAA-positive PD colon biopsies exhibited a seeding profile consistent with that previously reported in PD brain and cerebrospinal fluid samples(32, 37).

**Figure 2.**
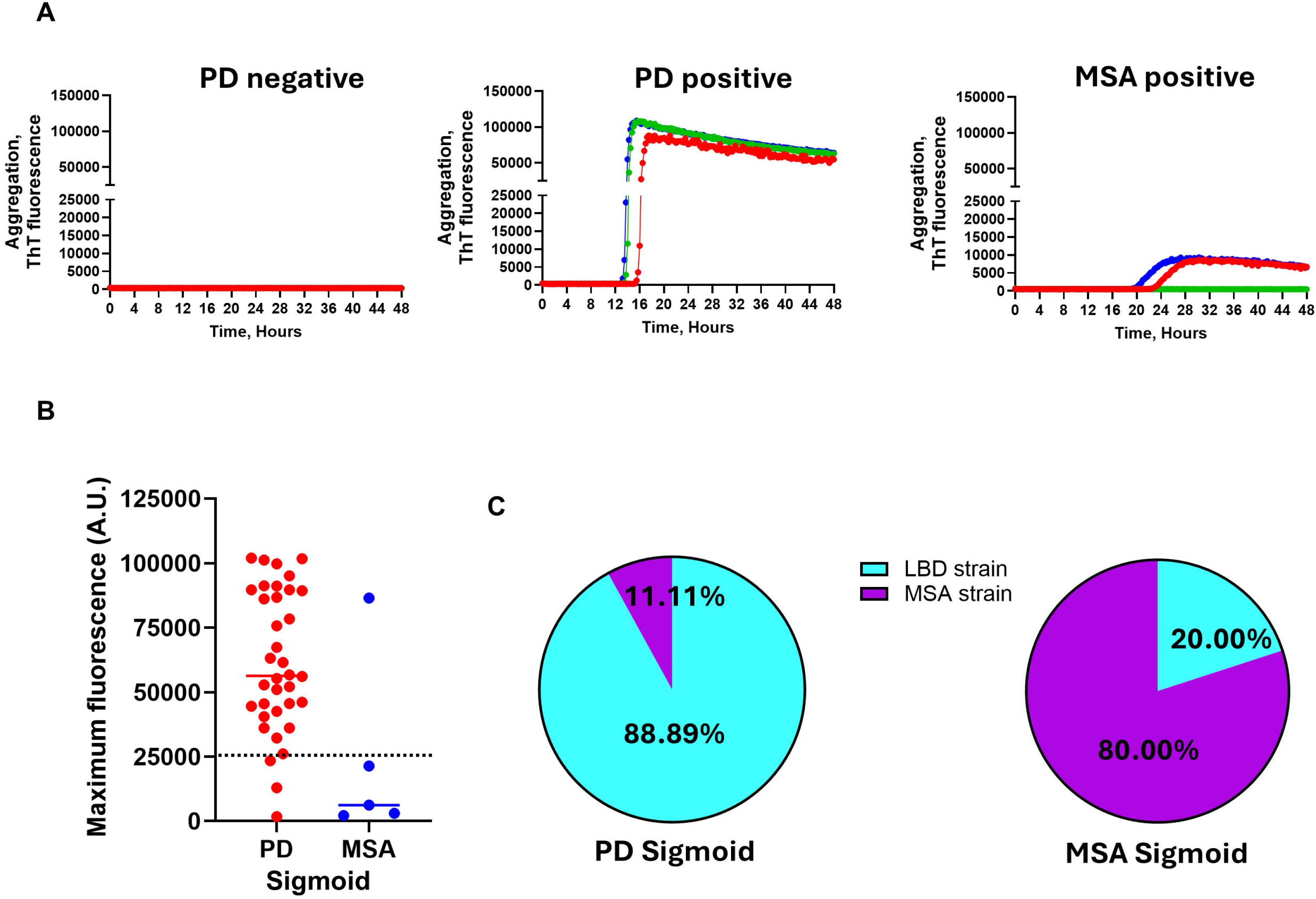
α-Synuclein Strain Typing in Sigmoid Biopsies. **(A)** Representative traces of ThT fluorescence during the α-synuclein SAA of sigmoid biopsies from Parkinson’s disease (PD) and multiple system atrophy (MSA) patients. Each line shows a different technical replicate. **(B)** Maximum fluorescence of the PD- and MSA-positive cases to identify the strain pattern in the two synucleinopathies. The data show the average fluorescence of the three technical replicates. The dotted line depicts the cutoff used to differentiate PD and MSA strains (25,000 fluorescence units). Each dot represents a different patient. Only positive cases are shown (36 PD, 5 MSA). **(C)** Percentage of PD and MSA samples in which the type-1 (LBD) or type-2 (MSA) strain was detected.

### Sigmoid SAA-Positivity Is Associated with PD Duration

The following PD analyses focus on the sigmoid mucosa, where SAA status varied according to PD severity. Because disease duration differed across the PD severity categories, we asked if disease duration rather than severity accounted for sigmoid SAA-positivity. Sigmoid SAA-positivity rose from 40.00% (10/25) in PD participants with 5 years of disease to 73.33% (11/15) at 5-10 years and 93.75% (15/16) 10 years (**Figure 1D**). Each 5-year increment in disease duration was associated with higher odds of sigmoid SAA-positivity (adjusted OR 2.82, 95% CI 1.39-5.75, p=0.004). Disease duration was also longer among sigmoid SAA-positive than SAA-negative PD participants (median of 9 versus 3 years, respectively; MWU: p=0.001).

Motor stage and motor burden showed numerically higher values among sigmoid SAA-positive PD participants, although neither association reached statistical significance. Sigmoid positivity was 13/16 (81.25%) at H&Y stages 3–5, which includes postural instability as a feature, versus 23/40 (57.50%) at stages 1-2 (Fisher’s exact p=0.127). MDS-UPDRS: Part III scores likewise did not differ between sigmoid SAA-positive and SAA-negative PD participants (median of 30.0 versus 26.5, respectively; MWU: p=0.43), and MDS-UPDRS: Part III was not associated with sigmoid SAA-positivity in logistic regression, either alone (OR 1.02 per point, 95% CI 0.97-1.07, p=0.52) or after adjustment for disease duration (OR 0.98, 95% CI 0.92-1.04, p=0.45), in which disease duration remained associated with SAA-positivity (OR 1.26, 95% CI 1.07-1.48, p=0.005). To separate disease duration from H&Y stage, we entered the data into a multivariable logistic regression model. Disease duration remained associated with sigmoid SAA-positivity (OR 1.237 per year, 95% CI 1.077-1.475, p=0.001), whereas H&Y stage did not (OR 1.027, 95% CI 0.181-6.107, p=0.976). Together, these analyses showed that sigmoid seeding competent α-syn aggregation is associated with disease duration rather than current motor symptoms or clinical stage.

Plasma NfL, a marker of axonal neurodegeneration(30), was elevated in PD relative to HC across all severity groups (KW: p<0.0001, **Table 1**, **Figure 1E**) but did not differ based on SAA-positivity status (two-way ANOVA: main effect of SAA status F_(1,50)_=0.832, p=0.366; severity x SAA interaction F_(2,50)_=0.134, p=0.874). The main effect of PD severity remained significant in this model (F_(2,50)_=3.32, p=0.0439, **Figure 1F**). Thus, plasma NfL parallels PD severity but does not track sigmoid SAA status, and disease duration remained the strongest independent correlate of sigmoid SAA-positivity.

#### Sigmoid SAA-Positivity is Not Accompanied by a Distinct Local Gut Phenotype

The gut-directed measures separated the PD severity groups but did not separate SAA-positive from SAA-negative PD participants. Bristol stool scores significantly decreased with increasing PD severity, reflecting harder stools and greater constipation (KW: p<0.0001, **Table 1**, **Supplementary Figure 1A**). However, stratifying by sigmoid SAA status did not impact Bristol Stool Score (two-way ANOVA: F_(1,44)_=1.502, p=0.2268; **Figure 3A**).

**Figure 3.**
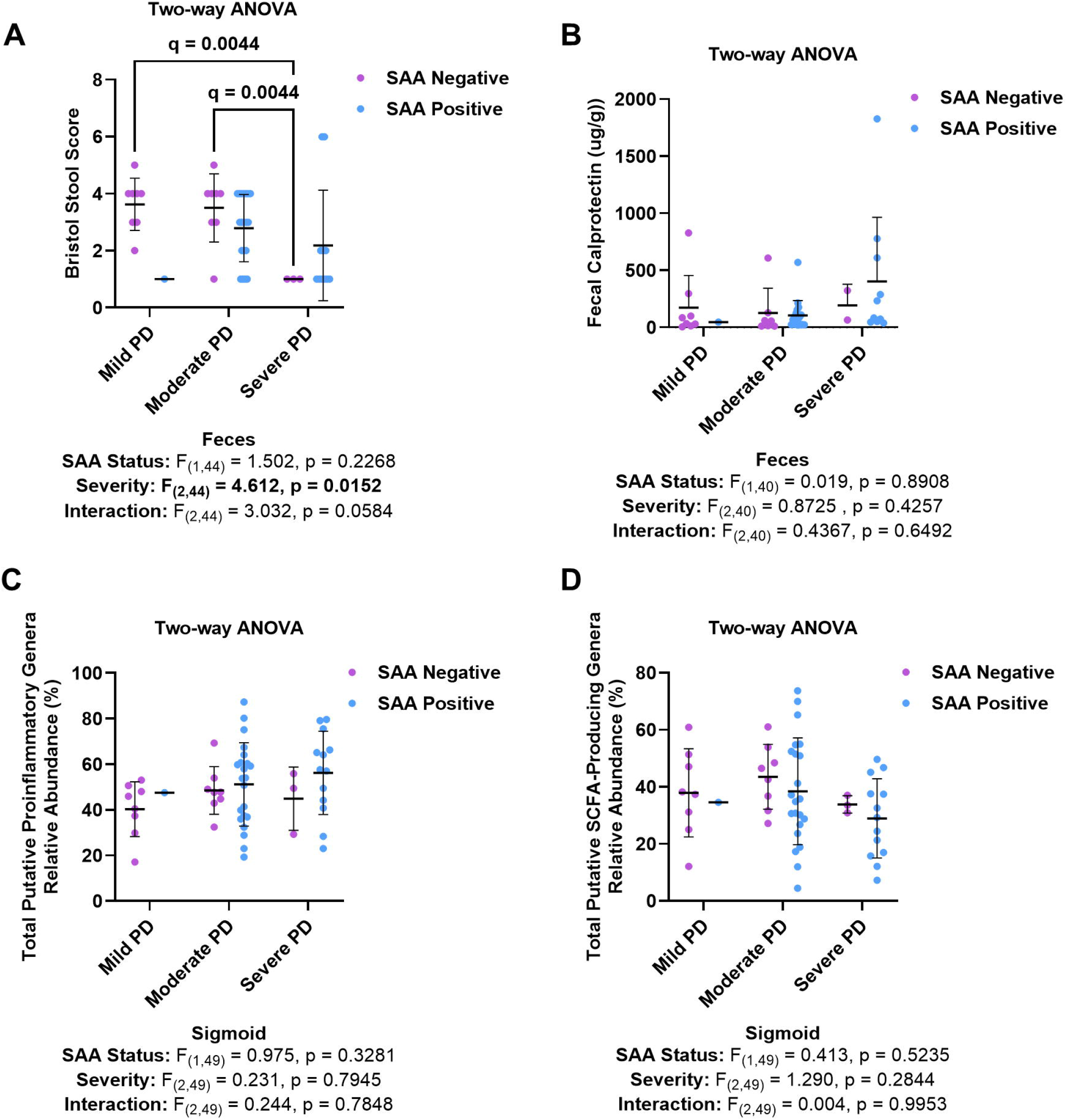
Gastrointestinal characteristics by sigmoid SAA status and PD severities. **(A)** Bristol Stool Form Scale scores, **(B)** stool calprotectin concentrations, **(C)** relative abundance of sigmoid putative pro-inflammatory bacterial genera, and **(D)** relative abundance of sigmoid putative short-chain fatty acid (SCFA)-producing genera. Within-PD two-way ANOVA evaluated severity, SAA status, and their interaction. Bristol score showed a PD severity effect (p=0.0152) but no statistically significant SAA effect (p=0.2268) or interaction (p=0.0584). No, statistically significant severity, SAA status or interaction effects were detected for the other measures. Indicated pairwise comparisons are BH-adjusted where applicable.

Stool calprotectin, a marker of intestinal inflammation(31), was statistically significantly higher in severe PD than in HC (q=0.0216, **Table 1**, **Supplementary Figure 1B**); however, calprotectin did not differ by sigmoid SAA status or PD severity, with no significant interaction (**Figure 3B**).

Compared to HC, PD sigmoid mucosa associated microbiota community showed the expected dysbiotic profile, with a higher relative abundance of putative pro-inflammatory genera (ANOVA: F_(3,78)_=4.134, p=0.009) and a lower relative abundance of putative anti-inflammatory, SCFA-producing genera (ANOVA: F_(3,78)_=3.247, p=0.026) (**Table 1**, **Supplementary Figure 1C-D**). When stratified by sigmoid SAA status, neither pro-inflammatory pathobionts nor anti-inflammatory SCFA-producing genera differed by SAA status, and no significant interactions were observed (**Figure 3C-D**). Thus, despite more severe intestinal inflammation, and more disrupted mucosa-associated microbial community associated with disease severity, PD sigmoid SAA-positivity was not accompanied by a distinguishable local inflammatory or microbial phenotype in these analyses.

#### SAA outperforms IHC in detecting α-Syn aggregates in the intestinal mucosa

Proteinase K-resistant, pS129 IHC staining was interpreted as evidence of pathological α-syn aggregates. Using proteinase K–treated pS129 IHC (**Figure 4A**), α-syn aggregates were not detected in any of the 24 ileal PD biopsies (**Figure 4B**), whereas SAA-positivity was 48.65% (18/37) in ileal PD. In contrast, IHC detected α-syn aggregates in 26.32% (10/38) of sigmoid PD biopsies (**Figure 4C**), compared with SAA-positivity of 64.29% (36/56). All HC biopsies evaluated by IHC were negative at both sites (ileum, 0/17; sigmoid, 0/20). Sigmoid IHC-positivity was greater with more severe disease (0/1, 0% mild; 7/30, 23.33% moderate; 3/7, 42.86% severe PD), although the small, uneven subgroups do not support a robust severity comparison (**Table 4**).

**Figure 4.**
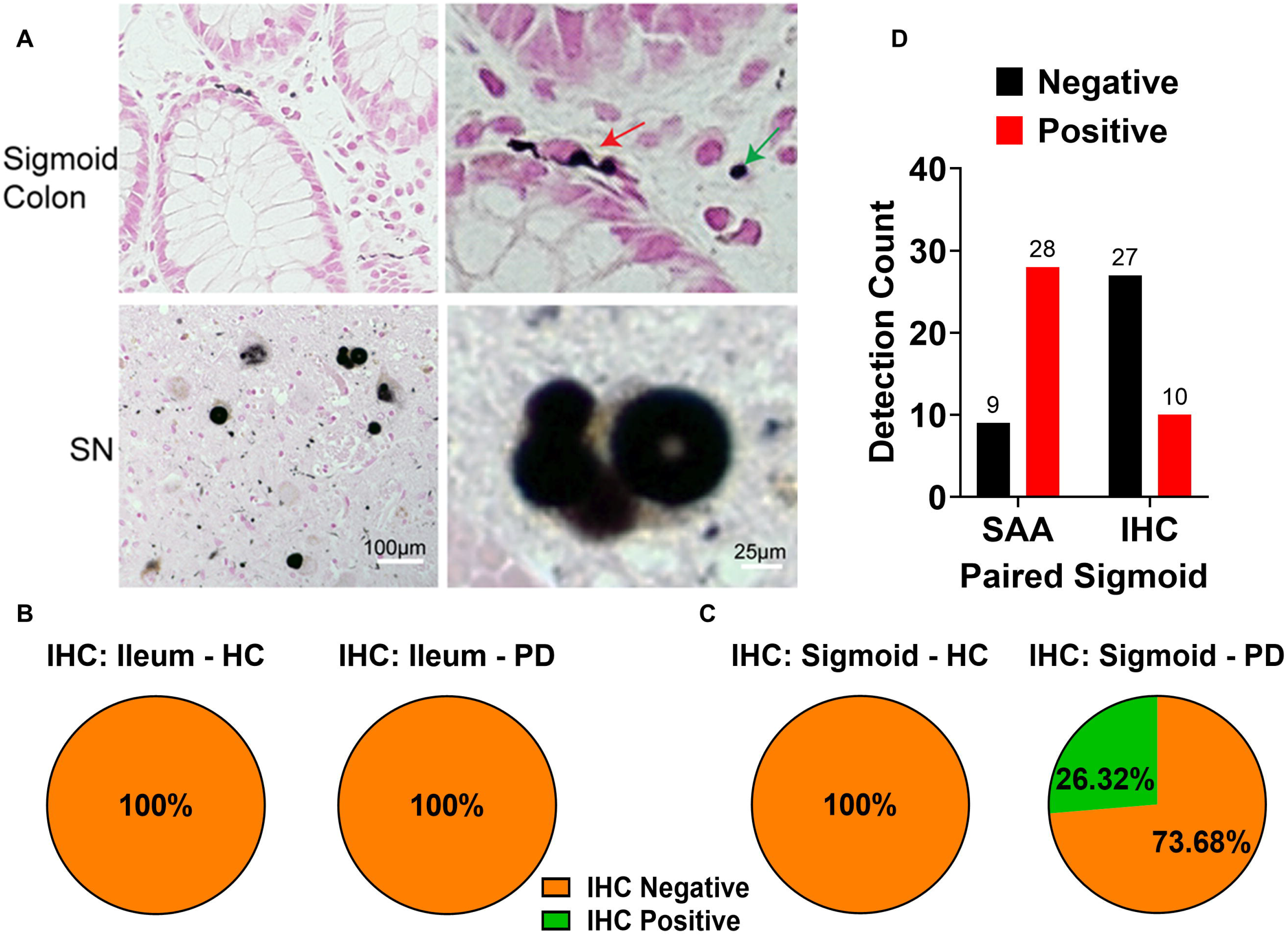
Comparison of intestinal α-synuclein immunohistochemistry (IHC) and seed amplification assay (SAA). **(A)** Representative proteinase K–treated phospho-Ser129 α-synuclein IHC in sigmoid mucosa and substantia nigra (SN) positive-control tissue; nickel-enhanced DAB appears black and nuclear fast red is the counterstain. Arrows indicate the structures interpreted as pS129-positive in the sigmoid specimen. Scale bars are as shown. **(B)** IHC was negative in all examined ileal biopsies (24 PD, 17 HC). **(C)** Sigmoid IHC was positive in 10/38 PD specimens and 0/20 HC specimens. **(D)** In 37 paired sigmoid PD specimens, SAA was positive in 28 (75.68%) and IHC in 10 (27.03%); 20 specimens were SAA-positive/IHC-negative and two were SAA-negative/IHC-positive (McNemar’s exact p=0.0001).

**Table 4.** Intestinal Immunohistochemistry (IHC) detection rates and diagnostic performance by biopsy site, participant group (HC, PD, MSA), and Parkinson’s disease severity.

| **Biopsy, Group** | **IHC Positive, n (%)** | **IHC Negative, n (%)** | **Total** | **Sensitivity (%)** | **Specificity (%)** | **PPV (%)** | **NPV (%)** |
|---|---|---|---|---|---|---|---|
| Ileum, HC | 0 (0%) | 17 (100%) | 17 | - | - | - | - |
| Ileum, PD | 0 (0%) | 24 (100%) | 24 | 0.00 | 100.00 | - | 41.46 |
| Sigmoid, HC | 0 (0%) | 20 (100%) | 20 | - | - | - | - |
| Sigmoid, PD | 10 (26.32%) | 28 (73.68%) | 38 | 26.32 | 100.00 | 100.00 | 41.67 |
| Sigmoid, MSA | 0 (0%) | 7 (100%) | 7 | 0.00 | 100.00 | - | 74.07 |
| **Biopsy, Severity** |  |  |  |  |  |  |  |
| Ileum, Mild PD | - | - | 0 | - | - | - | - |
| Ileum, Moderate PD | 0 (0%) | 19 (100%) | 19 | 0.00 | 100.00 | - | 47.22 |
| Ileum, Severe PD | 0 (0%) | 5 (100%) | 5 | 0.00 | 100.00 | - | 77.27 |
| Sigmoid, Mild PD | 0 (0%) | 1 (100%) | 1 | 0.00 | 100.00 | - | 95.24 |
| Sigmoid, Moderate PD | 7 (23.33%) | 23 (76.67%) | 30 | 23.33 | 100.00 | 100.00 | 46.51 |
| Sigmoid, Severe PD | 3 (42.86%) | 4 (57.14%) | 7 | 42.86 | 100.00 | 100.00 | 83.33 |

Clinical diagnosis was used as the reference standard. Diagnostic performance was calculated separately for PD versus HC and MSA versus HC using the corresponding biopsy-site controls. Severity-stratified PD comparisons reused the same site-specific HC group. Sensitivity, specificity, positive predictive value (PPV), and negative predictive value (NPV) were calculated from the observed counts. PPV and NPV reflect the proportions of cases and controls within each comparison and may not generalize to populations with different disease prevalence. The sigmoid MSA analysis was exploratory because of the small sample size. Dashes (–) indicate not applicable. HC, healthy controls; PD, Parkinson's disease; MSA, multiple system atrophy.

In 37 paired sigmoid biopsies, SAA detected α-syn aggregates at a 2.8-fold higher rate than IHC (75.68% [28/37] vs. 27.03% [10/37]; **Figure 4D**). The two assays were discordant in 22 participants: 20 were SAA-positive/IHC-negative and two were SAA-negative/IHC-positive (McNemar’s exact p=0.0001, **Data Source**). The paired difference in positivity was 48.65%. Diagnostic performance by site is summarized in **Table 5**. SAA sensitivity was 48.65% in ileum and 64.29% in sigmoid biopsies, with specificity of 100% and 97.50%, and negative predictive value of 65.45% and 66.10%, respectively. In the small duodenal set, SAA demonstrated 71.43% sensitivity with 100% specificity (PPV 100%, NPV 33.33%). IHC sensitivity was 0% in ileum and 26.32% in sigmoid biopsies, with 100% specificity at both sites (**Table 5**). In MSA, SAA detected α-syn aggregates in all five MSA-P sigmoid biopsies, whereas IHC was negative in all seven MSA biopsies (five MSA-P and two MSA-C).

**Table 5.** Diagnostic performance of α-synuclein seed-amplification-assay (SAA) and immunohistochemistry (IHC) in intestinal biopsies from Parkinson’s disease (PD), multiple system atrophy (MSA), and healthy control (HC) participants.

| **Assay** | **Sample Site** | **Comparison** | **Case (n)** | **HC (n)** | **Sensitivity** | **Specificity** | **PPV** | **NPV** |
|---|---|---|---|---|---|---|---|---|
| SAA | Ileum | PD vs HC | 37 | 36 | 48.65% | 100.00% | 100.00% | 65.45% |
| SAA | Sigmoid | PD vs HC | 56 | 40 | 64.29% | 97.50% | 97.30% | 66.10% |
| SAA | Duodenum* | PD vs HC | 7 | 1 | 71.43% | 100.00% | 100.00% | 33.33% |
| SAA | Sigmoid* | MSA vs HC | 7 | 40 | 71.43% | 97.50% | 83.33% | 95.12% |
| IHC | Ileum | PD vs HC | 24 | 17 | 0.00% | 100.00% | - | 41.46% |
| IHC | Sigmoid | PD vs HC | 38 | 20 | 26.32% | 100.00% | 100.00% | 41.67% |
| IHC | Sigmoid* | MSA vs HC | 7 | 20 | 0.00% | 100.00% | - | 74.07% |

Clinical diagnosis was used as the reference standard. Diagnostic performance was calculated separately for PD versus HC and MSA versus HC at each biopsy site. Cases (n) represent the PD or MSA group specified in each comparison. Sensitivity, specificity, positive predictive value (PPV), and negative predictive value (NPV) were calculated from the observed counts. PPV and NPV reflect the study-specific proportions of cases and controls and may not generalize to populations with different disease prevalence. \*Duodenal and MSA analyses were considered exploratory because of their small sample sizes. The same site-specific HC groups were reused for corresponding PD and MSA comparisons. A dash (—) indicates that PPV was not estimable because no positive IHC results were observed. PD, Parkinson's disease; MSA, multiple system atrophy; HC, healthy controls.

## Discussion

Four key PD and MSA findings emerge from this study. First, seeding-competent α-syn aggregates can be detected in mucosal biopsies with high observed specificity against HC (97.50-100%), approaching that reported for CSF SAA(3), although no direct comparison was performed. Sensitivity varied by GI region: 48.65% ileum and 64.29% sigmoid. Duodenum sampling was exploratory, with 71.43% (5/7) testing SAA-positive, but only one HC biopsy was available, precluding reliable specificity estimation. Single biopsy sampling may have contributed to false-negative results; whether analyzing additional biopsies would improve sensitivity requires further investigation.

Second, seeding-competent α-syn was also detected in sigmoid biopsies from MSA-P, extending reports in CSF and brain(35, 40). Intestinal α-syn seeding is therefore not exclusive to PD. Most SAA-positive PD sigmoid exhibited an LBD-associated Type 1 seeding profile (88.89%), whereas most SAA-positive MSA biopsies exhibited an MSA-associated Type 2 profile (80.00%). These findings are consistent with disease-associated seeding profiles previously reported in CSF and skin(32,37). Our results suggest that the α-syn strain is retained in intestinal biopsies, suggesting that SAA may serve to differentiate patients affected from PD from MSA.

Third, the clinical correlates of sigmoid seeding-competent α-syn separated into two clusters. Disease duration was the only variable significantly associated with sigmoid SAA-positivity in the models tested, whereas MDS-UPDRS: Part III scores and plasma NfL were higher in participants with more severe disease without distinguishing SAA-positive from SAA-negative participants. The relationship was region-dependent: sigmoid SAA-positivity increased with disease duration and severity, whereas ileum SAA-positivity remained stable. Constipation, stool calprotectin, and sigmoid mucosal dysbiosis differed with PD severity, but likewise did not distinguish SAA-positive from SAA-negative PD participants. However, our proof-of-concept, cross-sectional study, with relatively small sample size, does not necessarily establish independence of SAA α-syn aggregates from the PD local GI environment.

Fourth, SAA detected α-syn pathology at a higher rate than proteinase K-resistant pS129 IHC in the in the sigmoid mucosa of the same PD and MSA participants, and in the ileum, SAA was positive in subset of PD patients despite uniformly negative IHC.

The association of sigmoid SAA-positivity with PD duration is an important finding. Duration remained associated with positivity in models incorporating H&Y stage or MDS-UPDRS: Part III scores, whereas motor severity measures were not independently associated. This suggests that the probability of detecting mucosal seeding activity may relate more closely to time since clinical diagnosis than to motor impairment. The higher positivity rate in longer-duration cases may also reflect more widespread intestinal pathology at this stage, and a single biopsy may therefore be more likely to detect α-syn seeding-competent aggregates, whereas multiple biopsies across the colon might be able to detect α-syn aggregates in early-stage disease. However, the cross-sectional design does not establish within-person accumulation of α-syn pathology, whether SAA predicts progression, or whether residual confounding explains part of the association.

No, statistically significant association with sigmoid SAA status was detected for Bristol stool scores, stool calprotectin, or the two microbiota measures. These features differed across HC and PD subgroups, but the within-PD analyses were limited by sample size and uneven SAA-positive and SAA-negative subgroup counts. The absence of statistically significant results should not necessarily be interpreted as evidence that mucosal seeding activity is biologically independent of intestinal inflammation, motility, or microbiome.

The anatomical pattern also warrants further study. Sigmoid SAA-positivity increased across disease-duration categories, whereas ileal SAA-positivity showed no clear trend in the available subgroups. Differences in participant composition, biopsy mass, sampled tissue layers, and regional seed distribution could all contribute. Binary positivity indicates detectable seeding activity rather than aggregate abundance. The present data cannot establish that ileal pathology precedes sigmoid pathology, infer a direction of gut–brain spread, or identify an optimal prodromal sampling site. Paired within-participant analyses and longitudinal sampling are needed to directly answer these questions.

The head-to-head comparison provides direct evidence of higher SAA-positivity than IHC in the sigmoid samples tested by both methods. Among 37 paired sigmoid biopsies, 28 (75.68%) were SAA-positive and 10 (27.03%) were IHC-positive; 20 were SAA-positive/IHC-negative and two were SAA-negative/IHC-positive. In contrast, IHC was negative in all 24 PD and 17 HC ileal biopsies examined. A potential explanation: (1) homogenate amplification can detect sparse seeding activity that may be missed in thin histological sections, (2) seeding-competent aggregates detected by SAA may correspond to small oligomeric species not detectable by standard IHC methods, and (3) the two assays interrogate different biological properties; SAA detects seeding competence, whereas IHC identifies selected immunoreactive structures. Thus, these data show higher SAA-positivity in a clinically diagnosed PD cohort, but not necessarily superior diagnostic specificity or interchangeability of the assays.

Our exploratory MSA findings complement prior evidence of intestinal abnormalities. We previously reported disrupted sigmoid mucosal ZO-1 tight junctions, increased TLR4 expression, and microbial pro-inflammatory dysbiosis in MSA(41). The seven MSA participants had relatively advanced motor impairment, comparable to our severe PD group but with shorter disease duration, as is typical of MSA as an often aggressive degenerative condition (**Table 2**). Detection of seeding-competent α-syn in MSA (5/7) sigmoid biopsies supports intestinal involvement and indicates that intestinal SAA-positivity is not specific to PD. Among the five SAA-positive biopsies, four exhibited an MSA-associated Type-2 seeding profile and one exhibited an LBD-associated Type-1 profile(42), suggesting heterogeneity. Positivity was only observed in the five MSA-P participants and neither of two MSA-C participants; this small, disease-duration-confounded comparison cannot establish subtype specificity but warrants further study.

These results expand upon previous reports demonstrating α-syn aggregate seeding detection in GI tissue(14–17). Reported outcomes vary by biopsy GI region, assay protocol, and cohort. Fenyi et al. examined biopsies from the antrum, sigmoid colon, and rectum(14), Shin et al. examined gastric biopsies in early-stage diagnosed PD(15), Vascellari et al. detected seeding activity in 22 of 23 PD duodenal biopsies(16), and Emmi et al. evaluated gastroduodenal and skin biopsies(17). These differences argue against treating GI SAA as a single test with uniform performance. The present cohort includes ileal and sigmoid biopsies, an exploratory PD duodenal and MSA sigmoid biopsies, paired PD sigmoid SAA–IHC comparisons, and analyses of disease duration and clinical features. It did not compare intestinal SAA and CSF or skin assays, so relative diagnostic performance cannot be established.

Important limitations include uneven samples at different GI sites and clinical stages, particularly for mild PD, duodenal biopsies, MSA subtypes, and IHC. Site-specific estimates were based on partially overlapping participants, and an independent validation cohort was not available. Study participants had clinically established PD or were HC rather than patients undergoing prospective assessment for parkinsonism. Observed specificity against HC does not establish differential diagnostic accuracy against MSA or other synucleinopathies. Because sensitivity and sample-based negative predictive value were limited, a negative intestinal SAA result cannot exclude PD. The cross-sectional design does not establish causality, prodromal sensitivity, or within-participant progression.

In addition to the scientific and diagnostic power of SAA-positivity, we emphasize the practical patient-oriented advantage to colonic tissue evaluations. Standard care of patients in the age group of PD or MSA diagnosis includes periodic cancer-screening colonoscopy. Adding colonic biopsies carries minimal risk, and the SAA assessment is readily available. With the new focus on early diagnosis, prodromal identification, and the potentials for neuroprotective therapies very early in PD, having a test that can be applied in public health at the community levels offers new horizons for patient identification. If our preliminary finding of regional differences between SAA detection in PD and MSA-P hold true in larger studies, colonic biomarkers may prove particularly useful for early diagnostic distinction because the two disorders are difficult to separate. Although we are not at that point, future efforts planning for such developments are pertinent to the present. Clearly, mucosal biopsies collected during clinical endoscopy provide access to peripheral α-syn pathology without a dedicated procedure (skin biopsy or CSF collection), and SAA testing is substantially more informative than IHC. These data provide a clinically pertinent research path and a design specification for future studies: biopsy region, stratification by disease duration, and direct benchmarking against CSF and skin assays in the same participants. Longitudinal cohorts spanning prodromal and early PD will determine if intestinal SAA adds information beyond established clinical measures.

## Supporting information

GTDB226 Classifier

## Data Sharing

Reference **Source Data** file. Microbiome data are available in the NCBI BioProject database under accession number PRJNA1511883. The classifier script used for data analysis is provided as a Supplementary Document. Additional data is available on request to corresponding author.

## Acknowledgements

The authors gratefully acknowledge the support of: Kristi Lawrence, Ankur Naqib, Neha Joshi, Corey Neumann, and Michelle Villanueva.

## Authors’ Roles

A.K. conceived and designed the study, performed sigmoidoscopy and endoscopy procedures, collected mucosal biopsies, and co-wrote the first draft and then edited the manuscript. P.A.E. performed data analysis and co-wrote the first draft of the manuscript; C.S. supervised the seed amplification assay measurements, designed the data analysis, interpreted the data, and edited the manuscript; R.M.V designed the study, interpreted the data, and edited the manuscript; M.P. performed seed amplification assay measurements, designed the data analysis, interpreted the data, and edited the manuscript; S.P. performed seed amplification assay measurements, designed the data analysis, interpreted the data, and edited the manuscript; B.K. performed immunohistochemistry staining and scoring, interpreted the data and edited the manuscript; M.S. interpreted the data, and edited the manuscript; J.D. performed immunohistochemistry staining and scoring, interpreted the data and edited the manuscript; M.B. performed immunohistochemistry scoring, interpreted the data and edited the manuscript; M.A.M. interpreted the data, and edited the manuscript; A.S. designed the study, interpreted the data, and edited the manuscript. C.G.G. designed the study, interpreted the data, and edited the manuscript. All authors reviewed and approved the final manuscript and have access to the data and the study design.

## Declarations

### Ethics approval and consent to participate

The study protocol (ORA: 23102401) was reviewed and approved by the Rush University Medical Center Institutional Review Board (IRB), Chicago, Illinois, USA. All participants provided written informed consent prior to enrollment and participation in the study. All procedures were conducted in accordance with institutional guidelines and the ethical standards of the Declaration of Helsinki.

### Consent for publication

All authors have reviewed and approved the last version of the manuscript and consent to its publication.

### Patient and Public Involvement

Patients and members of the public were not involved in developing the research questions, selecting outcome measures, designing or conducting the study, or preparing this manuscript. They were not involved in developing plans for dissemination of the findings.

---

**Supplementary Figure 1.**
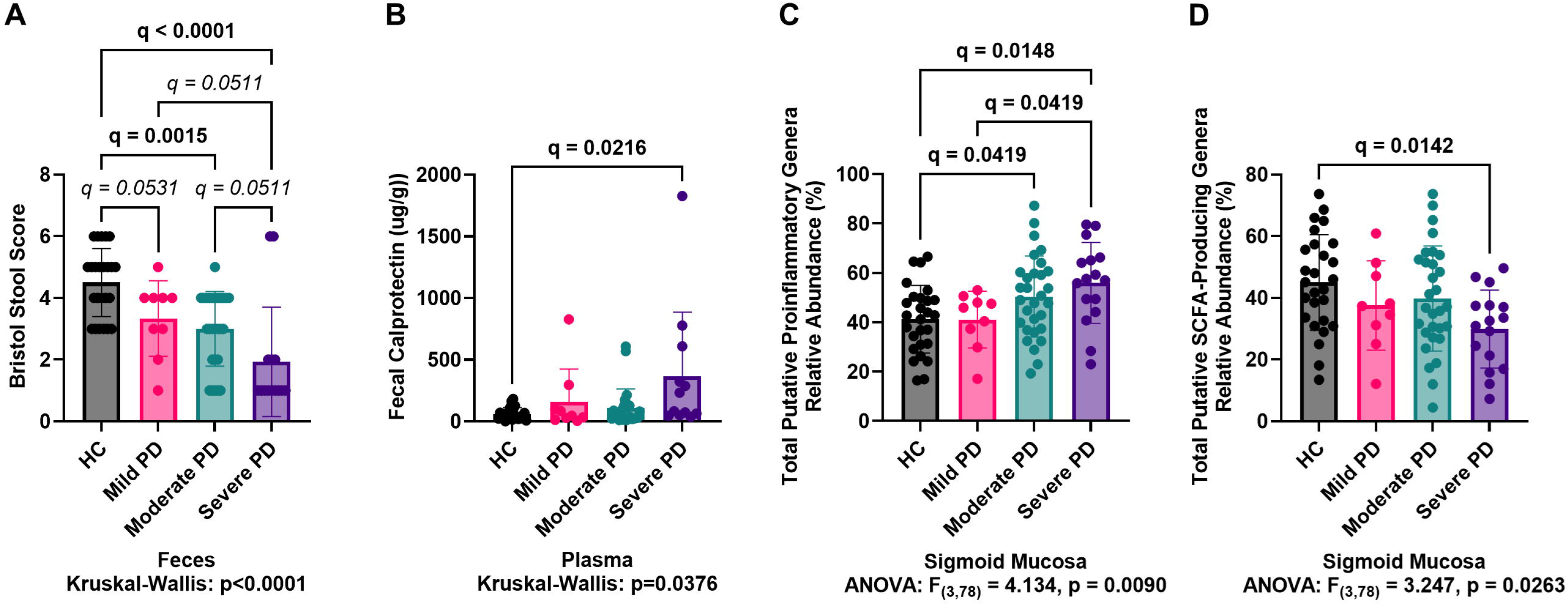
Gastrointestinal characteristics across PD subgroups. **(A)** Bristol Stool Form Scale scores across HC (n=40) and mild (n=9), moderate (n=31), and severe (n=16) PD groups (Kruskal-Wallis p<0.0001). **(B)** Stool calprotectin concentrations differed overall (Kruskal–Wallis p=0.0376). **(C)** Relative abundance of sigmoid putative pro-inflammatory bacterial genera differed across groups (ANOVA p=0.0090). **(D)** Relative abundance of sigmoid putative short-chain fatty acid (SCFA)-producing genera differed across groups (ANOVA p=0.0263). Indicated post hoc comparisons are reported as BH-adjusted q-values.

