## Supplemental Methods for "Seed Amplification Assay–Based Detection of Alpha-Synuclein Aggregates Outperforms Immunohistochemical Analysis in the Intestinal Mucosal Biopsies of Parkinson’s Disease and Multiple System Atrophy Participants"

**Supplementary Material**

**Supplementary Methods**

***Recombinant α-Synuclein Preparation***

Recombinant human α-Syn was produced and purified using an established protocol with modifications(29, 30). Briefly, C-terminal 6xHis-tagged human αSyn was expressed in BL21(DE3) pLysS E. coli cells following induction with 0.1 mM IPTG. After bacterial harvest, cells were suspended in lysis buffer containing 50 mM NaH₂PO₄ (pH 8.0), 300 mM NaCl, 10 mM imidazole, 1 mM PMSF, 0.1 mM TCEP, and 1 mg/ml lysozyme and disrupted by intermittent probe sonication. Cellular debris was removed by centrifugation at 12,000 x g for 15 min at 4 °C, followed by ultracentrifugation at 100,000 x g for 30 min at 4 °C. The clarified supernatant was subjected to nickel-affinity chromatography using Immobilized Metal Affinity Chromatography (IMAC). Recombinant α-Syn was recovered by elution with 250 mM imidazole, and α-Syn-containing fractions were subsequently dialyzed against PBS. Preformed aggregates were removed using a 100-kDa molecular-weight cutoff filter. Purified protein was pooled, aliquoted, and stored for subsequent SAA experiments.

***Immunohistochemistry***

Formalin-fixed paraffin-embedded biopsies were deparaffinized, rehydrated, and treated with proteinase K (40 μg/mL) for 20 min at 37 °C. Following peroxidase inactivation (0.3% hydrogen peroxide and 0.1% sodium azide, 30 min) and blocking for 1 hour with 2% bovine serum albumin with 5% goat serum diluted in buffer (20 mM Tris-HCl (pH 7.4), 150 mM NaCl, and 0.05% Triton X-100), sections were incubated overnight at 4 °C with an antibody against α-Syn phosphorylated at serine 129 (EP1536Y, Abcam, 1:50,000) in blocking buffer(1). Signal was detected using a biotinylated anti-rabbit antibody, ABC-HRP, and nickel-enhanced DAB(2, 3). Sections were counterstained with nuclear fast red (VWR). Two to four slides per biopsy were scored by three reviewers blinded to the study groups. Reviewers scored slides on a 0–3 scale (0=no signal, 1=mild, 2=moderate, 3=frequent staining) (4).

Conventional intestinal α-syn IHC has shown limited diagnostic specificity for PD, potentially reflecting physiological immunoreactivity and the rarity of pathological inclusions. In contrast, protease-resistant PS129 is a more robust approach for detecting pathological α-Syn aggregates and has demonstrated promising results in colon biopsies (4).
